# WIMOAD: Weighted Integration of Multi-Omics Data with Meta Learning for Alzheimer’s Disease Diagnosis

**DOI:** 10.1101/2024.09.25.614862

**Authors:** Hanyu Xiao, Jieqiong Wang, Shibiao Wan

**Affiliations:** Department of Genetics, Cell Biology and Anatomy, University of Nebraska Medical Center, Omaha, NE, United States, 68198; Department of Neurological Sciences, University of Nebraska Medical Center, Omaha, NE, United States, 6819

**Keywords:** Alzheimer’s Disease, Multi-Omics, DNA Methylation, Meta Learning, Stacking, Ensemble Learning, Weighted Score Fusion

## Abstract

**INTRODUCTION:** Alzheimer’s disease (AD), the most prevalent subtype of dementia, is characterized by a gradual decline in brain cognitive function. Early detection is critical for initiating timely interventions that may delay the severe progression of the disease. Recent advances in next-generation sequencing (NGS) offer promising, non-invasive, and cost-effective strategies for AD screening. However, most current approaches rely on single-omics data, which may fail to capture the complex biological heterogeneity among individuals.

**METHODS:** We introduce WIMOAD, a stacking ensemble and weighted multi-omics integration for AD diagnosis. It leverages paired gene expression and methylation data from ADNI and presents a meta learning framework for multi-cognitive stage classification during AD progression.

**RESULTS AND DISCUSSION:** WIMOAD outperforms existing integration methods in AD diagnosis, effectively capturing complex multi-omics patterns linked to clinical outcomes. Its interpretability also facilitates the detection of novel biomarkers across different omics layers.

## 1. BACKGROUND

Alzheimer’s disease (AD) is the most common subtype of dementia, characterized by a progressive decline in cognitive functions, notably in memory, thinking, and reasoning ^1^. It is closely associated with aging and exerts a persistent impact on cognitive functions ^2^. For primary healthcare and disease screening, the ability to achieve early and efficient diagnosis of AD is crucial for effective intervention and treatment ^3^. Typically, AD is characterized by the A/T/N framework ^4^. The majority of research relies on phenotypic data, particularly brain imaging ^5,6^. However, idealized imaging data remains limited, and the neuropathological diagnosis using cerebrospinal fluid (CSF) is invasive and harmful to patients ^7^. As pathophysiological changes gradually accumulate in amino acids, metabolism, and neuroinflammation, newly registered patients show considerable heterogeneity in the impaired cognitive domains, which will lead to increasing diagnostic costs ^8,9^, underscoring the need for more precise and individualized diagnostic approaches ^10–12^.

With the progress in sequencing techniques, genetic data are increasingly being utilized as external validation of AD hallmarks as a less-expensive and less-invasive measurement ^13^. For example, researchers have identified many genetic risk factors for AD identified by Single Nucleotide Polymorphism (SNP) in Genome-Wide Association Studies (GWAS) ^14,15^. As one of the main components of the epigenetic data and highly correlated with aging ^16^, DNA methylation level is found to be increased in peripheral cells of AD patients while correlating with worse cognitive performances and *APOE* polymorphism ^17,18^. However, considering the complex etiology of AD, which is related to aging, neurodegeneration, and pathological pathways and more, relying on one data modality only may underestimate other related risk factors in this complicated process since one omics can not convey all the information needed.

Therefore, the integration of multiple types of genetic data will greatly enhance the effectiveness of current AD research. For computational diagnostic models, leveraging multi-omics data will improve their accuracy, reliability, and interpretability ^19–21^. Nevertheless, how to combine data from different omics layers to provide a holistic view of biological systems remains the major challenge of this field. One general solution is to summarize all results from transcriptomic, proteomics, metabolomics, etc., on brains and other tissues and form a comprehensive understanding of the impact of one gene alteration in individual clinical trajectories ^22–25^. Multi-Omics Factor Analysis (MOFA), which represents high-dimensional variables to a smaller number of latent factors, is also brought up in this field^26^. iCluster ^27^, JIVE ^28^, and SLIDE ^29^ are all commonly used tools. In AD studies, Bao et al. ^30^ proposed SBFA that incorporates imaging and biological data for functional assessment questionnaire (FAQ) score prediction. Various artificial intelligence (AI)-based algorithms, mostly for disease diagnosis, have also been introduced ^31–33^, but methods proposed for AD diagnosis are still rare.

Despite these advancements, significant gaps remain in integration studies with AI-based models. Firstly, there is less attention on methylation data, which is highly related to aging and AD ^34–36^. Secondly, widely used direct data concatenation ^37^ often fails to preserve omics-specific information, as each omics has distinct data structures and biological representations. To fill this gap, we present WIMOAD, which incorporates meta learning and ensemble learning blocks for multi-omics integration for AD diagnosis. Our design is inspired by and builds upon our previous studies, which successfully applied ensemble strategies for protein subcellular localization ^38,39^, and now extends to the more complex challenge in neurodegenerative disease. WIMOAD synergistically leverages specialized classifiers for patients’ paired gene expression and methylation data and performs classification under different cognitive stages. The resulting scores are then stacked as the new representation for training a meta model for each omics. The prediction results of two distinct meta models are integrated with optimized weights for the final decision-making, providing higher performance than using single omics only. Benchmarking tests demonstrated that WIMOAD outperformed state-of-the-art multi-omics integration methods. We also applied the SHAP explainer ^40^ and found the most contributing biomarker genes among different omics layers, highlighting WIMOAD’s interpretability for new biomarkers detection.

## 2. METHODS

### 2.1 Datasets

The data used in this paper were collected from the Alzheimer’s Disease Neuroimaging Initiative (ADNI) database (adni.loni.usc.edu). ADNI is a longitudinal multicenter study that collected clinical, imaging, genetic, and biochemical biomarkers for early detection and tracking of recruited cohorts across different time points. For our model, we collected the data of 591 people’s gene expression and methylation profiles as model input following the criteria that the genetic profiles from different omics are paired for a certain sample (Originally we had 744 gene expression profiles and 649 methylation data records. The rest of the samples which only has one omics data were eliminated). Among them, there are 203 Normal Controls (CN) subjects (age: 76.38 ± 6.51, F/M: 101/102), 180 Early Mild Cognitive Impairments (EMCI) subjects (age: 71.72 ± 7.29, F/M: 81/ 99), 113 Late Mild Cognitive Impairments (LMCI) subjects (age: 74.51 ± 8.27; F/M: 45/68), which is 293 Mild Cognitive Impairments (MCI) and 95 Alzheimer’s Diseases (AD) (age: 77.05 ± 7.90, F/M: 35/60). The demographic information of the data is displayed in **Table 1**. For subsequent binary group classification tasks, we have reprocessed the original categories as follows: all samples, excluding the AD group, were categorized into a “patient” (PT) group to facilitate ‘PT-AD’ binary classification. Furthermore, the EMCI and LMCI groups were combined into a single MCI group, enabling the execution of other binary classification tasks related to MCI. The genetic profiling data (gene expression and methylation) are from participants’ blood samples.

**Table. 1.** The demographic information of the Selected Participants. Data are mean ± standard deviation (std).

| Diagnosis | Samples | Age (mean±std) | Sex (F/M) |
| --- | --- | --- | --- |
| CN | N = 203 | 76.38 ± 6.51 | 101/102 |
| EMCI | N = 180 | 71.72 ± 7.29 | 81/ 99 |
| LMCI | N = 113 | 74.51 ± 8.27 | 45/68 |
| AD | N = 95 | 77.05 ± 7.90 | 35/60 |

*Abbreviation: CN: Normal Controls; EMCI: Early Mild Cognitive Impairments; LMCI: Late Mild Cognitive Impairments; MCI: Mild Cognitive Impairments; AD: Alzheimer’s Diseases; F: Female; M: Male.*

### 2.2 Overview of WIMOAD Framework

WIMOAD is a weighted score fusion model based on the stacking ensemble of multiple base classifiers. As illustrated in the pipeline (**Figure 1A)**, gene expression and methylation data were extracted and paired according to patient ID to serve as model inputs. The model extracted the most variable expressed genes and the most variable methylated genes as features for binary classification from gene expression (*Exp*) and methylation data (*Met*), separately. For each data type, several machine learning classifiers, noted as base models, were applied independently to create new representations of the original datasets with the prediction scores for meta models. Finally, the meta model prediction results of *Exp* and *Met* were combined using a weighted fusion mechanism. Subsequent optimization was performed for each classifier and the ensemble weight to enhance the integration model performance. The optimized ensembled result was used to make the final decision on AD diagnosis. The model was validated using Leave-One-Out Cross Validation (LOOCV) tests. For base models in this study, we tested Support Vector Machine (SVM) ^41^, Random Forest (RF) ^42^, Naïve Bayes (NB) ^43^, Logistic Regression (LR) ^44^ Multi-Layer Perceptron (MLP) ^45,46^, and K-Nearest Neighbors (KNN) ^47^. The meta models selected for training in the framework were LR, RF, Linear Regression, and MLP.

**Fig. 1.**
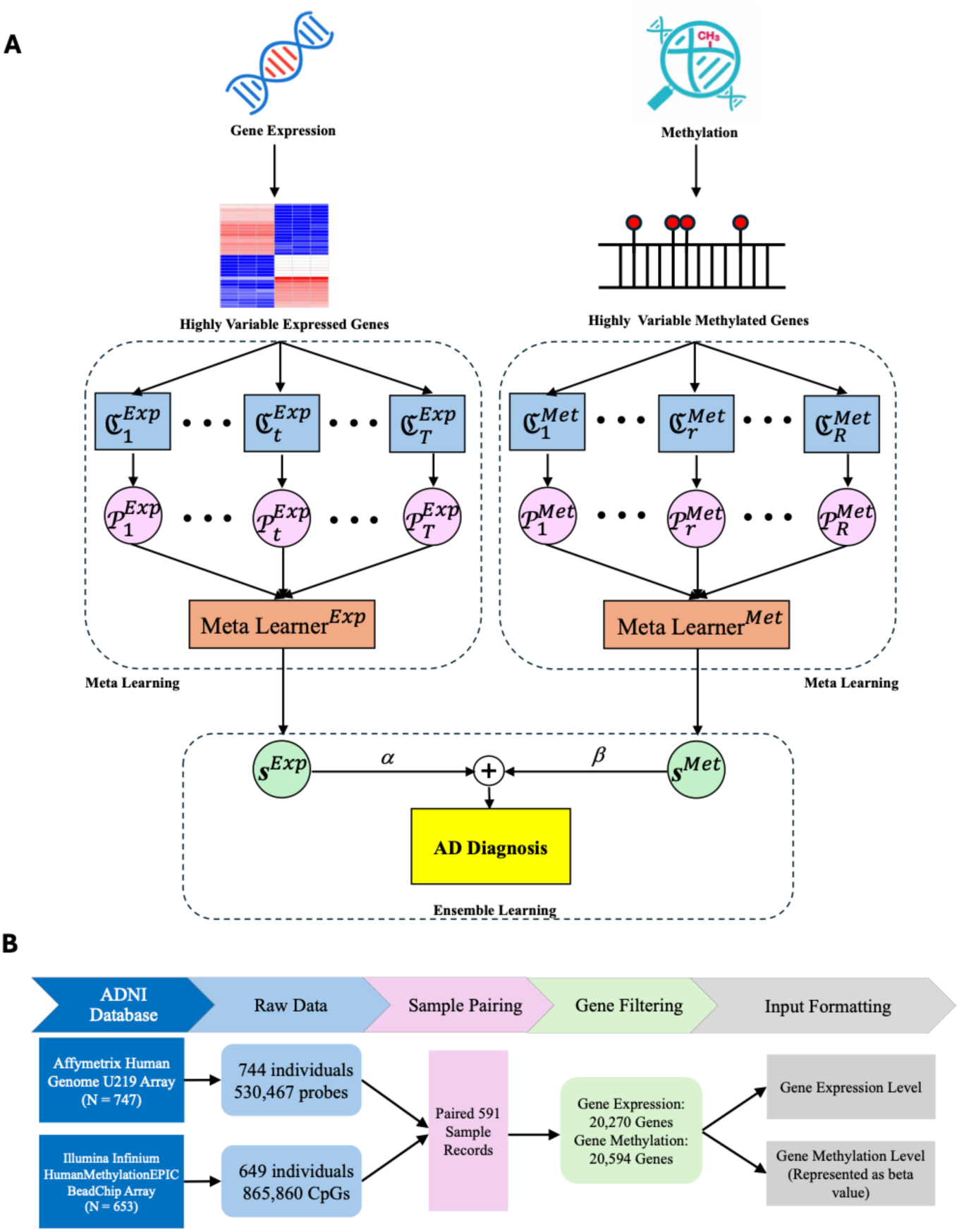
Data Processing and Model Contruction. (A)The Workflow of WIMOAD. The process begins by identifying the most variable features from paired gene expression and methylation data for classification. For each omics data, different classifiers (ℭ) were trained. The outputs of the basic classifiers (P) were considered as the new training sets for two distinct meta models, which used the predictions of base classifiers as inputs and generated the overall prediction scores (**s**). For multi-omics integration, each meta model is assigned a weight (α and β, β = 1-α) for ensemble learning, which also controls the contributions of each meta model to the final decision. **(B) Multi-omics data preprocessing.** Gene expression and methylation data were collected and preprocessed from ADNI. After sample pairing (591 samples selected), gene filtering, and input formatting, two matrices were yielded for further analysis.

### 2.3 Preprocessing of multi-omics data

The workflow of the multi-omics data preparation is illustrated in **Figure 1B**. Specifically, for gene expression, the Affymetrix Human Genome U219 Array is utilized for peripheral blood samples. The raw expression values generated by this platform were first normalized using the Robust Multi-chip Average (RMA) method, resulting in 530,467 probes corresponding to 49,293 transcripts from 744 samples. These probes were subsequently mapped and annotated according to the human genome reference (hg19). To ensure the specificity and reliability of gene-level expression, we selected the probe sets with high specificity that are uniquely designed for one transcript. Genes with missing information in the annotated data were excluded for further analysis. The filtered data contains 20,270 annotated genes.

For methylation, the Whole-genome DNA methylation profiling was conducted using the Illumina Infinium HumanMethylationEPIC BeadChip Array. The original data samples were normalized with the dasen method for downstream quality control (QC) including p-value criteria filtering, sex and sample ID verification, with 649 samples remaining ^48^. We obtained beta values for a total of 865,860 CpG sites by analyzing the channel signals. The identified CpG sites were also mapped and annotated with hg19, resulting in methylation data for 20,594 genes. For each gene, the methylation level was represented by the average beta value of its associated CpG sites. Through sample paring, 591 samples in total with both gene expression and methylation data were finalized during sample paring for later model establishment.

### 2.4 Feature Selection

For a supervised learning model, in the case of the “curse of dimensionality” and to enhance prediction efficiency while simultaneously reducing the consumption of computational resources, feature selection is a key process for model prediction. For each binary classification group, we applied ANOVA to evaluate the variability of gene expression/gene methylation within the group. Genes were ranked in descending order based on their F-value ^49^, with higher values indicating greater within-group variance. The top 1000 genes that show significant variance were selected for further analysis. The selection is done separately in two omics. The F-value is calculated by the ‘SelectKBest’ package in scikit-learn with ‘f_classif’ function. The ANOVA selection is only applied in the training sets.

### 2.5 Meta Learning

Ensemble learning leverages the diversity of individual classifiers to achieve improved overall performance of the computational model. Except for general voting, stacking is also commonly used, which combines the predictions of base-level classifiers together to establish the meta level dataset for decision-making and is found to outperform voting ^50^.

Within each omics-based classification task, multiple classifiers (base models) were employed to obtain first-level predictions. These predictions were subsequently stacked and utilized as new representations for training a meta learner. The meta learner’s predictions served as the final output, effectively integrating results from various base models to enhance predictive performance. In this study, we implemented two stacking strategies: (1) stacking only the base model predictions, and (2) stacking both the base model predictions and selected features as a comprehensive representation of the data. The key distinction of these strategies lies in whether the meta learner utilizes the model predictions only or incorporates the selected features from the input data.

### 2.6 Ensemble Learning

In omics integration research, a common approach is to concatenate different types of data directly before classification. However, in this study, *Exp* and *Met* data exhibit substantial differences in their data distributions and feature characteristics, which led to poor classification performance. Consequently, we applied prediction score fusion to construct an integration model for multi-omics data. Initially, a meta model was independently trained for each omics through meta learning mentioned above. The predictions from meta learners were weighted combined to derive the final prediction score of WIMOAD, which was calculated as:

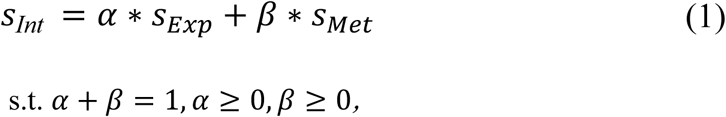

where *s*_Int_ is the integrated prediction score, *s*_*Exp*_is the score generated by the *Exp* meta model and *s*_*Met*_is the score generated by the *Met* meta model. The *α* and *β* are parameters controling the weights assigned to different omics predictions. These coefficients are determined through screening from *α* = 0 to *α* =1 with an increment of 0.01 in the linear combination. When *α > 0.5, Exp* predictions have larger contributions to the ensemble learning; when *β > 0.5, Met* predictions have larger contributions to the ensemble learning; when *α = β = 0.5,* both omics are equally weighted during integration.

### 2.6 Evaluation of the Model Performance

We used Leave-One-Out Cross Validation (LOOCV) ^51^ to evaluate WIMOAD performance. Specifically, we measured accuracy (Acc), precision (Prec), Recall (Rec), F1-Score (F1), Matthews correlation coefficient (MCC), Specificity (Sp), G-measure (G), Jaccard Index (Jacc) and Area Under Curve (AUC):

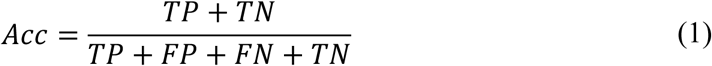

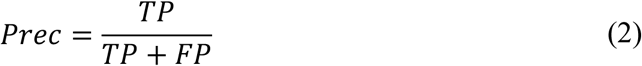

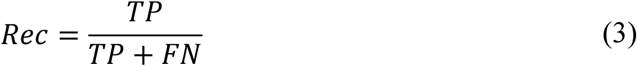

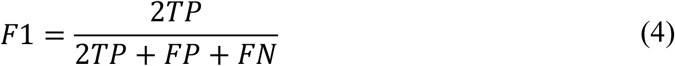

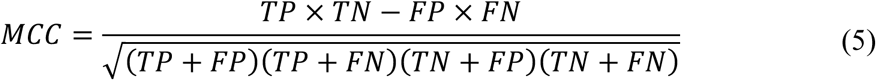

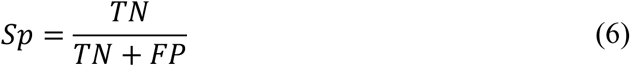

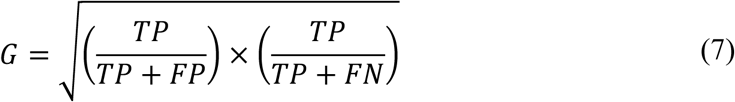

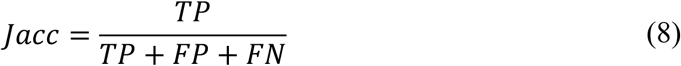

Among these, Acc measures the overall correctness of the predictions, Prec determines the reliability of positive predictions, when Rec evaluates the model’s ability to detect actual positive and Sp focuses on true negative cases. F1 balance the Prec and Rec, which is suitable for imbalanced cases. MCC, G and Jacc are for imbalance dataset results evaluation. AUC measures the area under the Receiver-Operating Characteristic (ROC) curve, evaluating the model’s ability to distinguish the two different classes across different thresholds. When performing random guessing, the AUC = 0.5.

### 2.7 Model Interpretation with SHAP

To make our model interpretable, we utilized the Kernel SHAP Explainer ^40,52^ for multi-kernel classifiers for different omics input. Given that different omics data modalities convey distinct types of information, interpreting each modality separately allows us to identify key genes contributing to the prediction results, providing a comprehensive understanding of the biological processes involved and highlighting critical genes that may be overlooked when considering a single data source. In addition, since we introduced the stacking strategy, multiple explainers were applied to different classifiers in each omics to see whether there are overlaps among the base models in contributing gene selection. In this process, genes were ranked by their absolute SHAP values, and the top 15 most influential genes were selected for each kernel explainer.

## 3. RESULTS

### 3.1 Meta Learning Enhances AD Diagnostic Performance Across Individual Omics Layers

To evaluate the impact of incorporating meta learning on the diagnostic performance of single omics data, we compared the AUC evaluation metrics of nine binary groups with base model and meta model prediction on *Exp* and *Met* data, visualized as ROC curves in **Figure 2** and **Figure S1**. Overall, introducing meta learning resulted in notable improvements in both omics. Particularly, there was about 2.35% to 16.91% abosolute improvement in the *Exp* AUC performance matrix and about 4.61% to 16.52% in the *Met* AUC after applying the meta learning strategy. Both the two stacking strategies achieved consistently better results than each single classifier in the binary group classification tasks. To be more specific, for base models, SVM and LR exhibited the most competitive performance across the two omics (SVM achieved highest in **Figure 2A-C, and Figure S1A, C, E, F, G, I**, while LR ranked first in **Figure 2D, F, and Figure S1H**. The two classifiers had the same performance in **Figure 2E, and Figure S1D**.). The AD vs. LMCI group demonstrated the most substantial improvement in the diagnosis rate after applying the meta learning strategy (**Figure 2A**) in *Exp* data, while the AD vs. LMCI reached the highest improvement for *Met*, respectively. The meta learning scheme in WIMOAD significantly improved the prediction power on single omics. The results also underscored the potential of meta learners to ensemble the complementary patterns from different base models and ultimately enhance the robustness and accuracy of the final predictions.

**Figure. 2.**
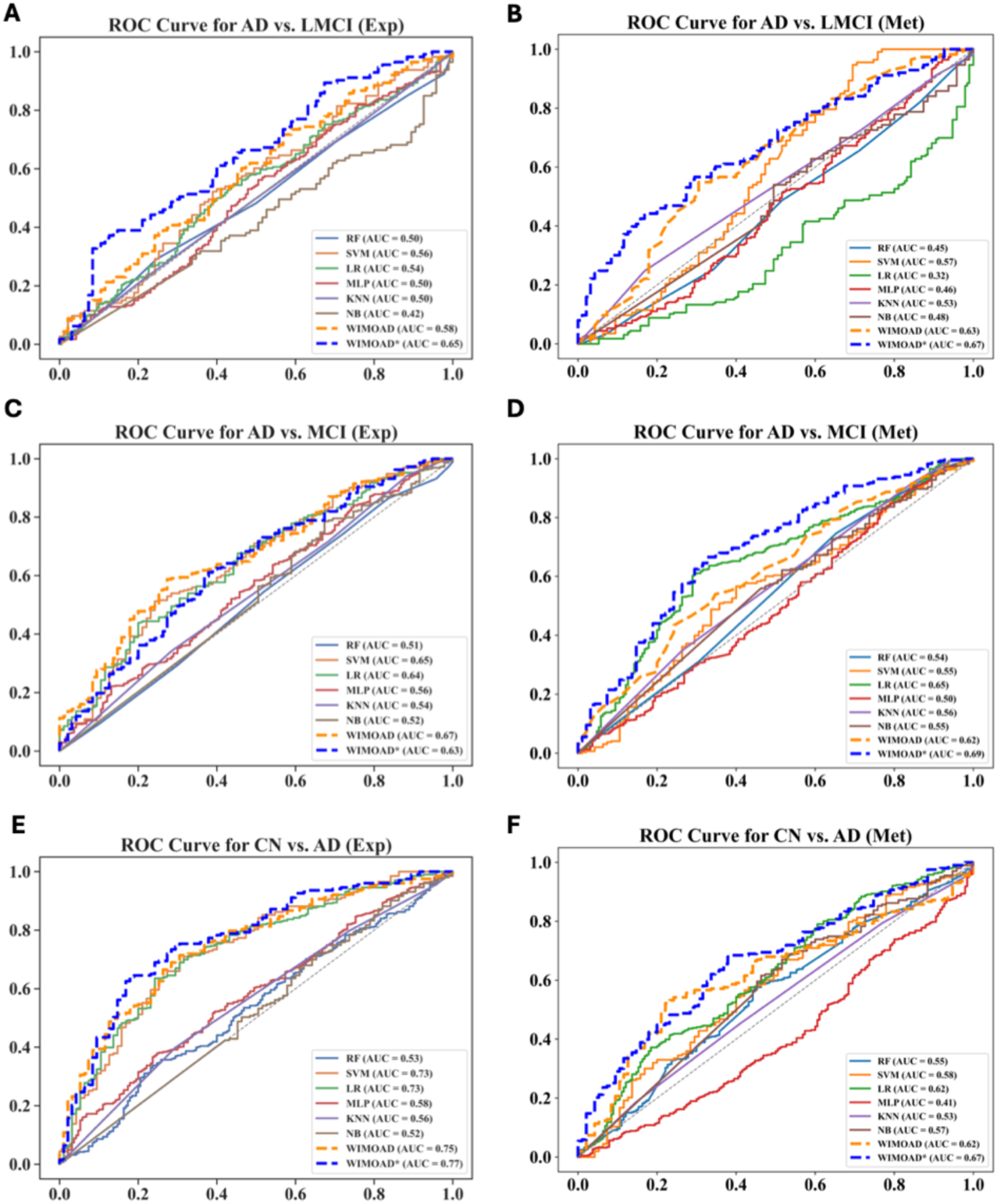
Comparing the Performance of Meta Learning Models Against Conventional Base Classifiers for AD diagnosis. All classifiers are trained with the same feature dimensions under LOOCV. The performance was measured using the metric introduced previously. **(A, C, E)** ROC curve on AUC comparison on gene expression data. (A) AD vs. LMCI group. (C) AD vs. MCI group. (E) CN vs. AD group. **(B, D, F)** ROC curve on AUC comparison on gene methylation data. (B) AD vs. LMCI group. (D) AD vs. MCI group. (F) CN vs. AD group. SVM: Support Vector Machine LR: Logistic Regression; MLP: Multilayer Perceptron; RF: Random Forest; NB: Naïve Bayes; KNN: K-Nearest Neighbor. WIMOAD*: Stacking strategy that adds the selected features for meta model training.

### 3.2 Base & Meta Model Selection for Meta Learning-Based AD Classification

To evaluate the impact of different model configurations, like hyperparameters of base model, the number of the base model, and the type of meta learners used in the framework, we analyzed WIMOAD performance under diverse conditions on two omics separately. All base models were subjected to hyperparameter optimization prior meta learning. A systematic comparison of the base model selected in this study quantities and their combinations, along with corresponding prediction results, was presented in **Figure 3**. The results indicated the advantage when using two to four base classifiers across all tasks. In contrast, selecting 5 to 6 classifiers or more was less competitive and did not consistently enhance predictive accuracy. For meta model training, *Exp* data with only base model predictions had the lowest variance across all conditions and higher AUC scores than adding original features (**Figure 3A-B**). In controversy, adding original features during meta model training for *Met* data yielded less variance and better performance compared to the best base models (**Figure 3C-D**). Furthermore, our investigation into stacking selected features with the predicted scores strategies suggested that the maximum AUC can be achieved through this strategy (e.g., AD vs. LMCI, CN vs. AD, CN vs. EMCI, CN vs. PT group in Exp, as shown in **Figure 3B** and AD vs. LMCI, AD vs. MCI, CN vs. AD, CN vs. EMCI group in Met, as shown in **Figure 3D**). In general, performing meta learning by stacking original features with base model predictions will greatly improve the model prediction and achieve performance than using base model predictions only under certain scenarios, but not applicable to all tasks.

**Figure. 3.**
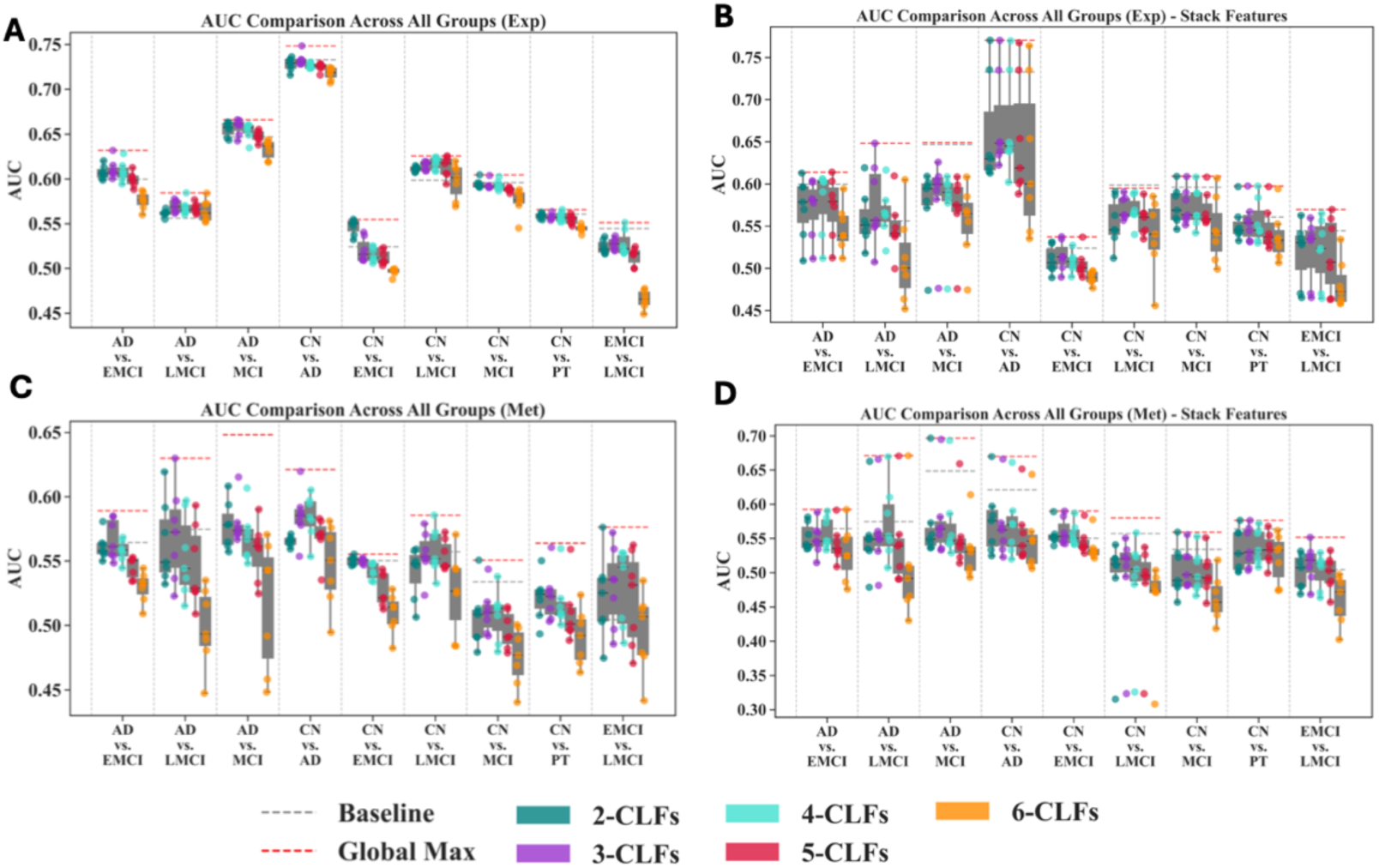
Impact of Base Model Settings on AD Diagnosis Performance Using Single Omics in Meta Learning. **(A, B)** WIMOAD improvement on *Exp* data compared to the best single classifier. (A) Meta learning with base model predictions only. (B) Meta learning with base model predictions and selected features. **(C, D)** WIMOAD improvement on *Met* data compared to the best single classifier, applied with stacking features strategy when training second-level meta models. (C) Meta learning with base model predictions only. (D) Meta learning with base model predictions and selected features. (2 - 6)-CLFs: Number of classifiers used as base models. Baseline: The highest AUC score of single classifier. Global Max: the highest AUC score across all combinations of classifiers and meta models in WIMOAD.

To further investigate the impact of meta model selection, we systematically compared the different meta learners and benchmarked their performance against commonly used averaging and voting methods under the same base model setting. Across all binary groups, training meta models for further prediction consistently performed better than simple voting schemes on both omics (**Figure 4 and Figure S2**). MLP and Linear Regression emerged as the two most effective meta models (noted as MLP1-4(M) and LR(M), respectively). MLP didn’t perform consistently better than the best base model yet achieved the highest at the AD vs. LMCI, AD vs. MCI group in *Exp* (**Figure 4A, C**). Linear Regression performed better than base models across all cases, when providing the best performance at the AD vs. LMCI, AD vs. MCI group in *Met* (**Figure 4B, D**), and CN vs. AD for both groups (**Figure 4E, F**). To add, both soft and hard voting either failed to surpass the best-performing base classifiers or had only slight improvement, highlighting the meta learning’s robustness for prediction power improvement in single omics for AD diagnosis.

**Figure. 4.**
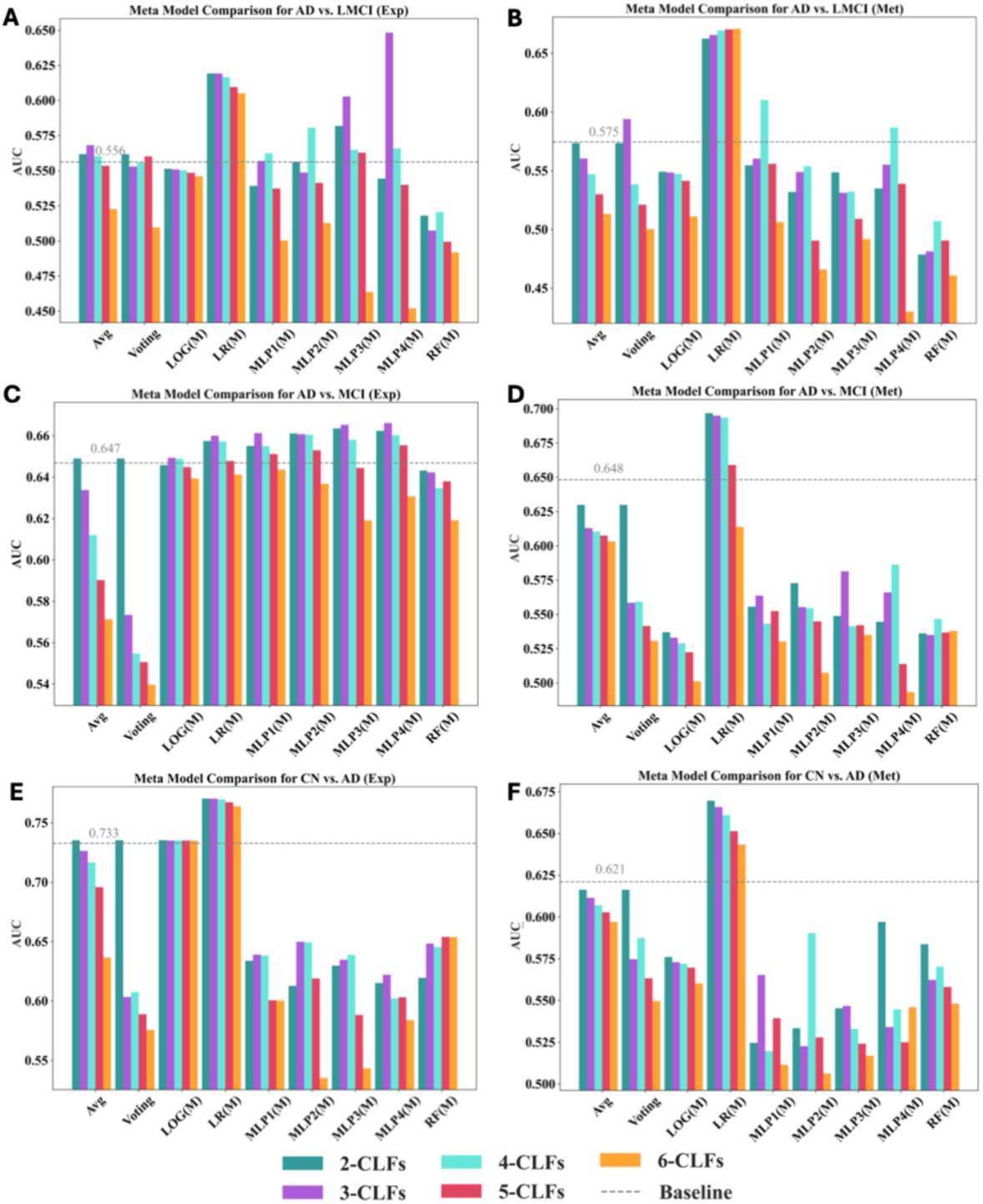
Impact of Meta Model Selection Versus Simple Averaging and Voting on AD Diagnostic Performance Using Fixed Base Model Combinations. **(A, C, E)** WIMOAD performance improvement using gene expression data only under different meta models. (A) AD vs. LMCI group. (C) AD vs. MCI group. (E) CN vs. AD group. **(B, D, F)** WIMOAD performance improvement using gene methylation data only under different meta models. (B) AD vs. LMCI group. (D) AD vs. MCI group. (F) CN vs. AD group. (2 - 6)-CLFs: Number of classifiers used as base models. Avg.: Average the prediction scores of base models for the ensemble; Voting: hard voting strategy for ensemble learning. LOG(M): Logistic Regression as the meta model for stacking ensemble; LR(M): Linear Regression as the meta model for stacking ensemble; MLP(1-4)(M): Multi-Layer Perceptron with different number of hidden layers as the meta model for stacking ensemble; RF(M): Random Forest as the meta model for stacking ensemble. Baseline: The highest AUC score of single classifier. Global Max: the highest AUC score across all combinations of classifiers and meta models in WIMOAD.

### 3.3 Integration Enhances Predictive Performance Beyond Individual Omics

The superiority of multi-omics integration over single omics approaches for AD diagnosis was assessed by a weighted ensemble of meta learner predictions from individual omics (*s_Exp_* and *s_Met_*) (**Figure 5)**. With optimized weights on *Exp* data, the value of feature integration and the potential for original sampling exceeded the performance of both *Exp* and *Met* meta model outputs. According to the AUC comparison, the integration model could outperform both omics when assigning weight from 0 to 1 when achieving peaks between 0.25 (**Figure 5H**) and 0.93 (**Figure 5A**). After applying meta learing, *Exp* was treated as the best model in 5 cases (**Figure 5A, D, F, G, H**) while *Met* for 4 cases (**Figure 5B, C, E, I**), the increase varies from 1.59% (AD vs. EMCI group, **Figure 5A**) to 8.96% (**AD vs. LMCI group, Figure 5B**) and 5.26% (EMCI vs. LMCI group, **Figure 5I**) to 19.40% (CN vs. AD group, **Figure 5D**) compare to the best and worst single model, respectively. We compared the prediction performance of the integrated model under the optimal weight condition to the best-performing single omics model (achieved under meta learning conditions) (**Figure S3**) and conducted McNemar’s Test (**Table S1)** to check whether there was a statistically significant difference, with the integration model demonstrating superior predictive power compared to the model constructed solely based on one omics. Compared to the original data without the meta learning module mentioned above, the overall improvement of WIMOAD was 4.03% to 31.29% for *Exp* data and 9.09% to 28.82% for *Met* data (**Table S2**). These results demonstrated that WIMOAD can effectively enhance the discriminative ability when mitigating the impact of weak-performing omics at the final integration stage via ensemble learning in multi-omics integration. All reported improvements are calculated as absolute differences.

**Figure 5.**
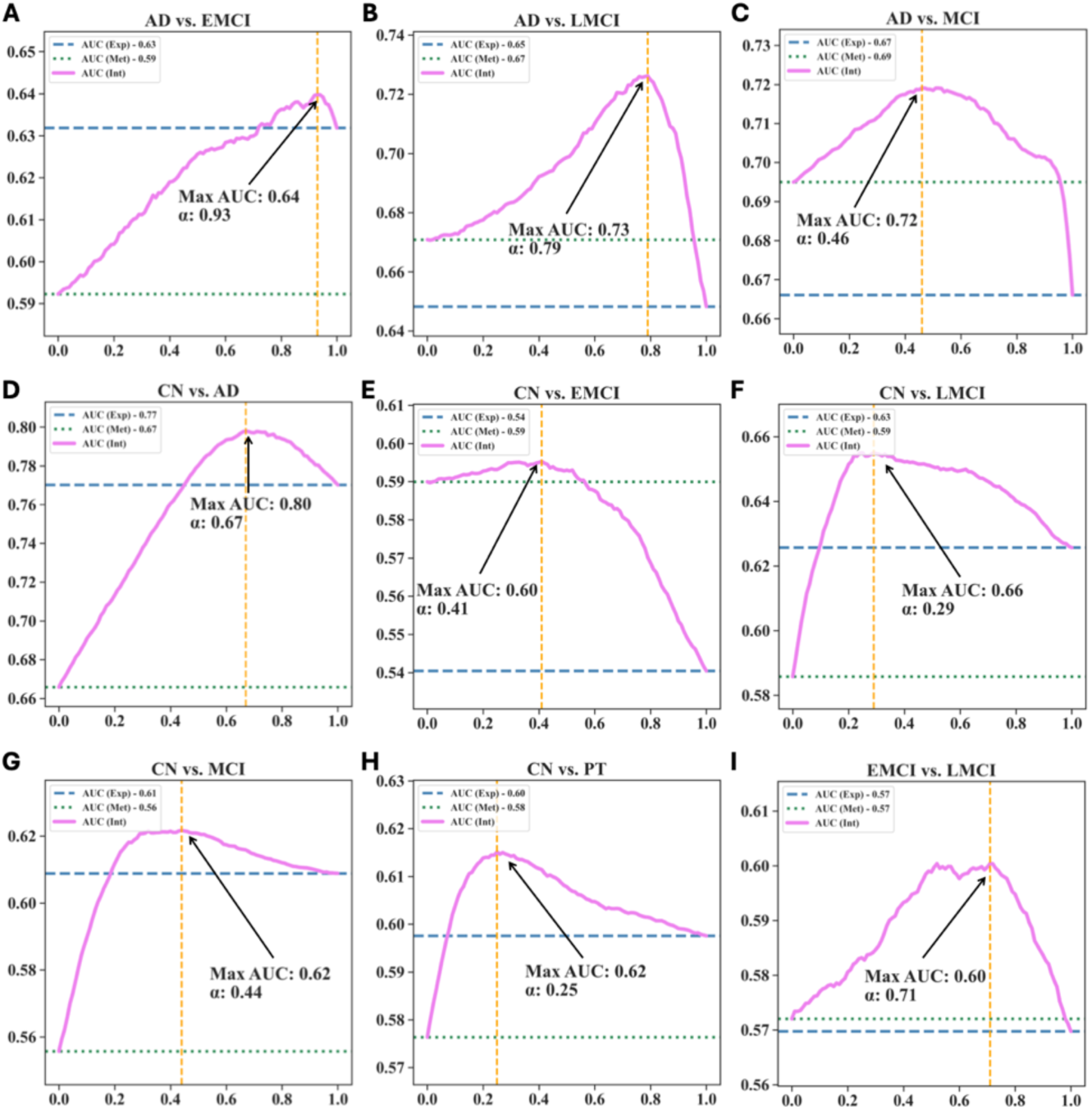
Enhanced Predictive Performance of the Integration Diagnostic Model Under Varying Weight Coefficients. The x-axis in each subfigure represents the increase of the integration coefficient α, which is the weight assigned to the prediction results of *Exp* meta model. The y-axis represents the accuracy of the model. The vertical dashed yellow line represents the highest AUC with respect to the weight coefficient α. (A) AD vs. EMCI group. (B)AD vs. LMCI group. (C)AD vs. MCI group. (D)CN vs. AD group. (E)CN vs. EMCI group. (F)CN vs. LMCI group. (G)CN vs. MCI group. (H)CN vs. PT group. (I)EMCI vs. LMCI group.

### 3.4 WIMOAD remarkably outperformed state-of-the-art predictors for AD diagnosis

To demonstrate the robustness and effectiveness of WIMOAD, we compared it with state-of-the-art predictive models with integration strategies using the paired ADNI data in this study (**Table. 2**). MOGLAM ^53^ is an end-to-end graph-based method for multi-omics integration. It applies a multi-omics attention mechanism to create new representative embeddings of different omics with weight, catching both shared and complementary information across different layers. IntegrationLearner, proposed by Mallick et al. ^54^, is a Bayesian ensemble method that combines information across several longitudinal and cross-sectional omics data layers and allows uncertainty quantification and interval estimation. LOOCV is applied to all methods for comparison.

**Table. 2.** Comparing state-of-the-art methods. All model apply ADNI data as input source. Avg: Average.

| Groups | WIMOAD |  |  | IntegrationLearner<sup>54</sup> |  |  | MOGLAM<sup>53</sup> |
| --- | --- | --- | --- | --- | --- | --- | --- |
|  | *Exp* | *Met* | *Int* | *Exp* | *Met* | *Int* | *Int* AUC |
|  | AUC | AUC | AUC | AUC | AUC | AUC |  |
| AD vs. EMCI | **0.63** | **0.59** | **0.64** | 0.52 | 0.53 | 0.50 | 0.34 |
| AD vs. LMCI | **0.65** | **0.67** | **0.73** | 0.44 | 0.45 | 0.44 | 0.39 |
| AD vs. MCI | **0.67** | **0.69** | **0.72** | 0.52 | 0.47 | 0.52 | 0.36 |
| CN vs. AD | **0.77** | **0.67** | **0.80** | 0.56 | 0.50 | 0.56 | 0.45 |
| CN vs. EMCI | **0.54** | **0.59** | **0.60** | 0.53 | 0.50 | 0.53 | 0.53 |
| CN vs. LMCI | **0.63** | **0.59** | **0.66** | 0.48 | 0.49 | 0.51 | 0.56 |
| CN vs. MCI | **0.61** | **0.56** | **0.62** | 0.44 | 0.49 | 0.47 | 0.54 |
| CN vs. PT | **0.60** | **0.58** | **0.62** | 0.478 | 0.52 | 0.52 | 0.53 |
| EMCI vs.<br>LMCI | **0.57** | **0.57** | **0.60** | 0.50 | 0.46 | 0.50 | 0.50 |
| Avg | **0.63** | **0.61** | **0.67** | 0.50 | 0.49 | 0.51 | 0.47 |

Across all the nine binary classification groups, the WIMOAD demonstrated consistently higher AUCs (0.67 on average compared to 0.51 using IntegrationLearner and 0.47 using MOGLAM), highlighting its capability to differentiate various AD stages at higher rates. Average all given tasks, WIMOAD achieved an AUC of 0.67 with integrated data, compared to 0.51 for IntegrationLearner and 0.47 for MOGLAM. Notably, in classification scenarios such as AD vs. LMCI and CN vs. AD group, WIMOAD reached AUCs of 0.73 and 0.80, respectively, outperforming the other two methods by substantial margins. These results highlighted the effectiveness of the WIMOAD framework.

### 3.5 WIMOAD is Interpretable to Identify Biomarkers for Different Cognitive Stages

To explore the biological relevance of WIMOAD’s predictions, we leveraged the SHAP explainer to interpret the model by quantifying the contribution of the highly variable expressed and methylated genes as inputs to the final model output at each omics layer. Take CN vs. AD for example, the top 15 genes with the greatest influence on the base model’s predictions were identified by ranking the genes according to the magnitude of their SHAP values, revealing those most critical to distinguishing between the two groups. (SVM, RF, NB, KNN, LR, MLP in **Figure 6A-F**, respectively). Remarkably, discernible variations emerged across different binary groups and omics data types. It became evident that the regulatory dynamics, manifested through gene upregulation or downregulation, yield bidirectional effects on the model’s decision boundaries, influencing the classification outcome for individual samples. After the intersection of the top 500 contributing genes among five classifiers, *TENM1 (Teneurin Transmembrane Protein 1)* was one of the genes present in the overlap, which is highly associated with AD diagnosis ^55–57^. Notably, this gene appeared as one of the top 15 contributing genes in SVM (**Figure 6A**), RF (**Figure 6B**), KNN (**Figure 6D**), and LR (**Figure 6E**), ranking the 1^st^ place in SVM and LR, and 6^th^ in RF and 8^th^ in KNN. The results demonstrated the biological interpretability of WIMOAD by identifying genes previously reported in the literature as established AD hallmarks, while also highlighting its potential to uncover novel biomarkers for AD screening.

**Fig. 6.**
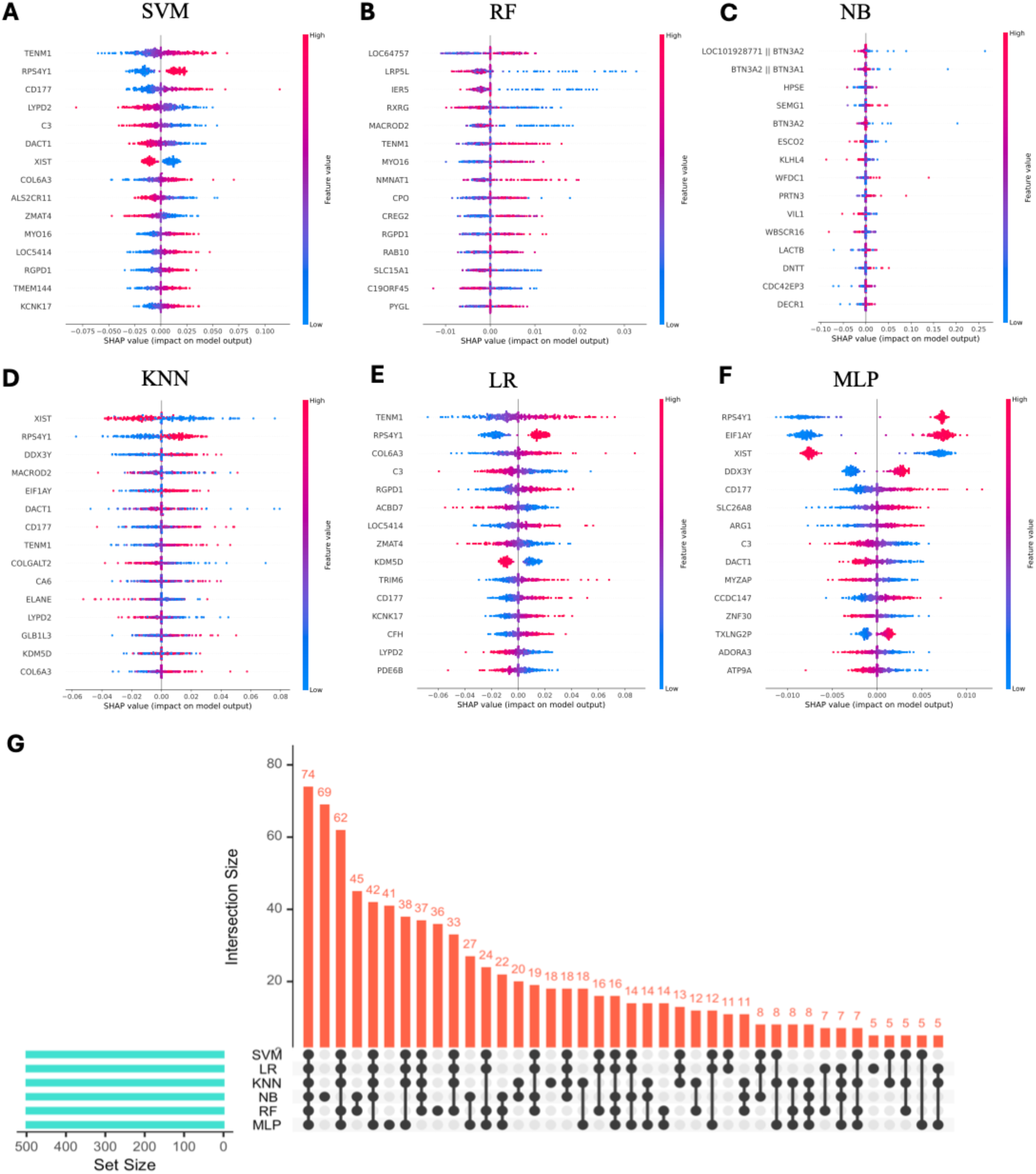
SHAP Plots for Model Explanation and Contributing Genes Detection. Top 15 most contributing genes and their influence on the model classification (sample being classified as AD) were exhibited. **(A-F)** SHAP summary plots for gene expression classifier of CN vs. AD group. (A) SVM. (B) RF: Random Forest. (C) NB: Naïve Bayes. (D) KNN: k-Nearest Neighbors (E) LR: Logistic Regression. (F) MLP:Multi-Lyaer Percenptron. The different colors illustrate the gene expression/methylation level of certain genes, and the SHAP values of the certain gene for each sample are denoted in the x-axis. Higher positive SHAP values for a certain gene represent the higher possibility that with the expression/methylation value, the model will classify the sample as AD and vice versa. **(G)** The Upset diagram summarized the number of overlapping genes of six gene sets generated from the top 500 contributing genes in each classifier.

## 4. DISCUSSION

In this study, we developed WIMOAD (Weighted Integration of Multi-Omics Data with Meta Learning for Alzheimer’s Disease Diagnosis), a stacking ensemble multi-omics integration framework for AD diagnosis. It incorporated gene expression and methylation data to distinguish different cognitive stages in AD accurately and effectively.

WIMOAD is a meta learning-based model that outperformed typical individual ML classifiers for AD diagnosis. As the convolutional and MLP-based classifiers and algorithms did not provide better performance with the datasets according to the single classifiers comparison, we established meta models that take the predictions from base classifiers as new training dataset, which is the new representative of the original data, to “learn” how to combine the results of different classifiers automatically for prediction performance improvement. After comparing to single classifiers and widely used averaging or voting techniques when leveraging multiple classifiers at the first layer, our meta models achieved consistently better performances across classification tasks in AD diagnosis at each omics level.

WIMOAD integrates multi-omics data by a late-integration scheme via a weighted ensemble. By assigning weights to the prediction scores generated by meta learners from each omics, the overall model performance was generally enhanced. This weighted fusion led to at least one weight setting where the integrated model surpasses any single-omics model. These results highlighted the effectiveness of WIMOAD in enhancing discriminative power by mitigating the influence of weak-performing omics through ensemble-based late integration. Moreover, this flexible framework enables the simultaneous incorporation of additional omics layers, making it possible to achieve higher diagnostic performance and offering a more comprehensive computational approach to AD screening. Notably, while proteomics profiles ^58^ were initially considered in the development of WIMOAD, data availability posed a significant constraint. After quality control and filtering, only 129 samples contained matched gene expression, methylation, and proteomics data, which were sufficient solely for the CN–LMCI binary classification task. Due to the limited availability of multi-omics samples, the current study focused on integrating gene expression and methylation data.

WIMOAD performs significantly better than state-of-the-art integration models for AD diagnosis. We tested other multimodal fusion models, such as IntegrationLearner from Mallick et al. ^54^, a novel Bayesian ensemble method that combines information across several longitudinal and cross-sectional omics data layers, and MoGLAM from Ouyang et al. ^53^, which integrates a dynamic graph convolutional network, attention mechanism, and omic integrated representation learning modules for fusing DNA methylation, miRNA, and mRNA expression profiles for disease classification. Comparative analysis revealed that WIMOAD consistently outperformed these methods across all classification groups. A likely reason for WIMOAD’s superior results is its use of meta learning and weighted score fusion to aggregate predictions from different classifiers, followed by decision-making, rather than directly concatenating data from various sources as input for predictions.

WIMOAD is biologically interpretable to characterize biomarkers genes for AD diagnosis. Instead of directly combining data, WIMOAD can extract specific representations from different data modalities simultaneously and fully use all the information for the prediction. By quantifying the contributions of the most variable genes separately, WIMOAD will contribute to the detection of new biomarkers in multi-omics for early diagnosis, biomarker discovery, and precision therapy design in AD studies. Given that the SVM model can currently only utilize KernelSHAP—an algorithm within SHAP with relatively high computational complexity and longer runtime—we have limited our presentation to the top ten genes (both expression and methylation) that most significantly influence the model’s predictions. Integrating SHAP into the decision-making process allows the visualization of how gene expression/methylation levels affect model predictions as well. For instance, a higher expression level of a particular gene correlates with a higher corresponding Shapley value, indicating that when the model detects high expression of this gene in a sample, it is more likely to classify the sample into a specific category. This demonstrates that the gene’s expression level has a direct impact on the model’s final prediction. Consequently, incorporating the SHAP explainer makes it feasible to identify new biomarkers. Additionally, in binary classification cases, the results obtained from different groups could potentially serve as markers for identifying the various stages in the progression from healthy (CN) to AD.

Despite the advantages, we admit that WIMOAD has some limitations. One limitation is that WIMOAD is currently designed for binary classification only. In other words, WIMOAD performs well at binary classification across different cognitive stages in AD (e.g., EMCI, LMCI). In addition, since the data all comes from the peripheral blood, the biomarker detection in the study needs further investigation about how it links with the change in the brain and how it will contribute to the mechanism of the AD process.

## 5. CONCLUSION

In this paper, we proposed a meta learning-based multi-omics integration model for Alzheimer’s Disease diagnosis named WIMOAD. It extracted information from both gene expression and paired methylation profiles of samples for model decision-making. To effectively integrate heterogeneous omics sources, WIMOAD adopted a stacking ensemble framework, where multiple base learners were trained independently on each omics, and their prediction outputs were treated as new training sets for meta learner that captured cross-modal predictive patterns. For the integration process, we implemented a weighted ensemble strategy in which each meta learner’s prediction was assigned an optimized weight at the final decision stage for prediction improvement. This late integration scheme not only improved robustness by reducing the influence of weak-performing modalities but also allowed flexible scalability to additional omics layers. Besides outperforming single classifiers for single omics, WIMOAD surpassed the most recent models presented that incorporate statistical analysis and deep learning algorithms. It was also interpretable through the SHAP explainer by quantifying top contributing genes at different omics layers for new biomarker detection during disease progression. One of the future directions for our research is to extend WIMOAD to be capable of performing multi-class classifications across different cognitive stages during AD progression and incorporating commonly utilized imaging data and the correlations across different data modalities to develop a more comprehensive multi-modality-based diagnostic model that enhances AD diagnostics’ robustness and clinical applicability in disease pathology.

## ACKNOWLEDGMENTS

HX developed the algorithm, conducted the experiments, analyzed the data, and implemented the WIMOAD package. JW co-supervised the whole project, edited and reviewed the manuscript. SW supervised the whole project, designed the study, and contributed to manuscript editing and review. All authors contributed to writing the manuscript and approved the final version. All authors participated in writing the paper. The manuscript was approved by all authors.

## DATA AVAILABILITY

All the data used in this manuscript are publicly available in the corresponding references. WIMOAD is available at https://github.com/wan-mlab/WIMOAD.

## Reference

1. Jack CR, Bennett DA, Blennow K, et al. NIA-AA Research Framework: Toward a biological definition of Alzheimer’s disease. Alzheimer’s &amp; Dementia. 2018;14(4):535–562. doi:10.1016/j.jalz.2018.02.018

2. Pini L, Pievani M, Bocchetta M, et al. Brain atrophy in Alzheimer’s Disease and aging. Ageing Research Reviews. 2016;30:25–48. doi:10.1016/j.arr.2016.01.002

3. Rasmussen J, Langerman H. Alzheimer’s Disease – Why We Need Early Diagnosis. DNND. 2019;Volume 9:123–130. doi:10.2147/DNND.S228939

4. Jack CR, Bennett DA, Blennow K, et al. A/T/N: An unbiased descriptive classification scheme for Alzheimer disease biomarkers. Neurology. 2016;87(5):539–547. doi:10.1212/WNL.0000000000002923

5. Bazarbekov I, Razaque A, Ipalakova M, Yoo J, Assipova Z, Almisreb A. A review of artificial intelligence methods for Alzheimer’s disease diagnosis: Insights from neuroimaging to sensor data analysis. Biomedical Signal Processing and Control. 2024;92:106023. doi:10.1016/j.bspc.2024.106023

6. Tong Y, Li Z, Huang H, Gao L, Xu M, Hu Z. Research of spatial context convolutional neural networks for early diagnosis of Alzheimer’s disease. J Supercomput. 2024;80(4):5279–5297. doi:10.1007/s11227-023-05655-9

7. Suganyadevi S, Pershiya AS, Balasamy K, Seethalakshmi V, Bala S, Arora K. Deep Learning Based Alzheimer Disease Diagnosis: A Comprehensive Review. SN COMPUT SCI. 2024;5(4):391. doi:10.1007/s42979-024-02743-2

8. Jack CR, Holtzman DM. Biomarker Modeling of Alzheimer’s Disease. Neuron. 2013;80(6):1347–1358. doi:10.1016/j.neuron.2013.12.003

9. Bateman Randall J., Xiong Chengjie, Benzinger Tammie L.S., et al. Clinical and Biomarker Changes in Dominantly Inherited Alzheimer’s Disease. New England Journal of Medicine. 2012;367(9):795–804. doi:10.1056/NEJMoa1202753

10. Friedland RP, Koss E, Haxby JV, et al. Alzheimer Disease: Clinical and Biological Heterogeneity. Ann Intern Med. 1988;109(4):298–311. doi:10.7326/0003-4819-109-4-298

11. Ferreira D, Nordberg A, Westman E. Biological subtypes of Alzheimer disease. Neurology. 2020;94(10):436–448. doi:10.1212/WNL.0000000000009058

12. Ringman JM, Goate A, Masters CL, et al. Genetic Heterogeneity in Alzheimer Disease and Implications for Treatment Strategies. Curr Neurol Neurosci Rep. 2014;14(11):499. doi:10.1007/s11910-014-0499-8

13. Hampel H, Lista S, Neri C, Vergallo A. Time for the systems-level integration of aging: Resilience enhancing strategies to prevent Alzheimer’s disease. Progress in Neurobiology. 2019;181:101662. doi:10.1016/j.pneurobio.2019.101662

14. Forte A, Lara S, Peña-Bautista C, Baquero M, Cháfer-Pericás C. New approach for early and specific Alzheimer disease diagnosis from different plasma biomarkers. Clinica Chimica Acta. 2024;556:117842. doi:10.1016/j.cca.2024.117842

15. Kamboh MI, Demirci FY, Wang X, et al. Genome-wide association study of Alzheimer’s disease. Transl Psychiatry. 2012;2(5):e117. doi:10.1038/tp.2012.45

16. Horvath S, Raj K. DNA methylation-based biomarkers and the epigenetic clock theory of ageing. Nat Rev Genet. 2018;19(6):371–384. doi:10.1038/s41576-018-0004-3

17. Di Francesco A, Arosio B, Falconi A, et al. Global changes in DNA methylation in Alzheimer’s disease peripheral blood mononuclear cells. *Brain*, Behavior, and Immunity. 2015;45:139–144. doi:10.1016/j.bbi.2014.11.002

18. Wei X, Zhang L, Zeng Y. DNA methylation in Alzheimer’s disease: In brain and peripheral blood. Mechanisms of Ageing and Development. 2020;191:111319. doi:10.1016/j.mad.2020.111319

19. Qiu S, Sun M, Xu Y, Hu Y. Integrating multi-omics data to reveal the effect of genetic variant rs6430538 on Alzheimer’s disease risk. Front Neurosci. 2024;18. doi:10.3389/fnins.2024.1277187

20. Das D, Ito J, Kadowaki T, Tsuda K. An interpretable machine learning model for diagnosis of Alzheimer’s disease. PeerJ. 2019;7:e6543. doi:10.7717/peerj.6543

21. Zhang J, Sun X, Jia X, et al. Integrative multi-omics analysis reveals the critical role of the PBXIP1 gene in Alzheimer’s disease. Aging Cell. 2024;23(2):e14044. doi:10.1111/acel.14044

22. Oka T, Matsuzawa Y, Tsuneyoshi M, Nakamura Y, Aoshima K, Tsugawa H. Multiomics analysis to explore blood metabolite biomarkers in an Alzheimer’s Disease Neuroimaging Initiative cohort. Sci Rep. 2024;14(1):6797. doi:10.1038/s41598-024-56837-1

23. Badhwar A, McFall GP, Sapkota S, et al. A multiomics approach to heterogeneity in Alzheimer’s disease: focused review and roadmap. Brain. 2020;143(5):1315–1331. doi:10.1093/brain/awz384

24. Nativio R, Lan Y, Donahue G, et al. An integrated multi-omics approach identifies epigenetic alterations associated with Alzheimer’s disease. Nat Genet. 2020;52(10):1024–1035. doi:10.1038/s41588-020-0696-0

25. Kodam P, Sai Swaroop R, Pradhan SS, Sivaramakrishnan V, Vadrevu R. Integrated multi-omics analysis of Alzheimer’s disease shows molecular signatures associated with disease progression and potential therapeutic targets. Sci Rep. 2023;13(1):3695. doi:10.1038/s41598-023-30892-6

26. Wang D, Gu J. Integrative clustering methods of multi-omics data for molecule-based cancer classifications. Quant Biol. 2016;4(1):58–67. doi:10.1007/s40484-016-0063-4

27. Shen R, Olshen AB, Ladanyi M. Integrative clustering of multiple genomic data types using a joint latent variable model with application to breast and lung cancer subtype analysis. Bioinformatics. 2009;25(22):2906–2912. doi:10.1093/bioinformatics/btp543

28. Lock EF, Hoadley KA, Marron JS, Nobel AB. Joint and individual variation explained (JIVE) for integrated analysis of multiple data types. Ann Appl Stat. 2013;7(1). doi:10.1214/12-AOAS597

29. Gaynanova I, Li G. Structural learning and integrative decomposition of multi-view data. Biometrics. 2019;75(4):1121–1132. doi:10.1111/biom.13108

30. Bao J, Chang C, Zhang Q, et al. Integrative analysis of multi-omics and imaging data with incorporation of biological information via structural Bayesian factor analysis. Briefings in Bioinformatics. 2023;24(2):bbad073. doi:10.1093/bib/bbad073

31. Mahendran N, Vincent P M DR. Deep belief network-based approach for detecting Alzheimer’s disease using the multi-omics data. Computational and Structural Biotechnology Journal. 2023;21:1651–1660. doi:10.1016/j.csbj.2023.02.021

32. Stahlschmidt SR, Ulfenborg B, Synnergren J. Multimodal deep learning for biomedical data fusion: a review. Briefings in Bioinformatics. 2022;23(2):bbab569. doi:10.1093/bib/bbab569

33. Lu P, Hu L, Mitelpunkt A, Bhatnagar S, Lu L, Liang H. A hierarchical attention-based multimodal fusion framework for predicting the progression of Alzheimer’s disease. Biomedical Signal Processing and Control. 2024;88:105669. doi:10.1016/j.bspc.2023.105669

34. Bollati V, Galimberti D, Pergoli L, et al. DNA methylation in repetitive elements and Alzheimer disease. Brain, Behavior, and Immunity. 2011;25(6):1078–1083. doi:10.1016/j.bbi.2011.01.017

35. Barrachina M, Ferrer I. DNA Methylation of Alzheimer Disease and Tauopathy-Related Genes in Postmortem Brain. Journal of Neuropathology & Experimental Neurology. 2009;68(8):880–891. doi:10.1097/NEN.0b013e3181af2e46

36. Scarpa S, Cavallaro RA, D’Anselmi F, Fuso A. Gene silencing through methylation: An epigenetic intervention on Alzheimer disease. Journal of Alzheimer’s Disease. 2006;9(4):407–414. doi:10.3233/JAD-2006-9406

37. Pammi M, Aghaeepour N, Neu J. Multiomics, artificial intelligence, and precision medicine in perinatology. Pediatr Res. 2023;93(2):308–315. doi:10.1038/s41390-022-02181-x

38. Wan S, Mak MW, Kung SY. Transductive Learning for Multi-Label Protein Subchloroplast Localization Prediction. IEEE/ACM Trans Comput Biol and Bioinf. 2017;14(1):212–224. doi:10.1109/TCBB.2016.2527657

39. Wan S, Mak MW, Kung SY. Ensemble Linear Neighborhood Propagation for Predicting Subchloroplast Localization of Multi-Location Proteins. J Proteome Res. 2016;15(12):4755–4762. doi:10.1021/acs.jproteome.6b00686

40. Lundberg SM, Lee SI. A Unified Approach to Interpreting Model Predictions.

41. Cortes C, Vapnik V. Support-vector networks. Mach Learn. 1995;20(3):273–297. doi:10.1007/BF00994018

42. Ho TK. Random decision forests. In: Proceedings of 3rd International Conference on Document Analysis and Recognition. Vol 1.; 1995:278–282 vol.1. doi:10.1109/ICDAR.1995.598994

43. Webb GI. Naïve Bayes. In: Sammut C, Webb GI, eds. Encyclopedia of Machine Learning. Springer US; 2010:713–714. doi:10.1007/978-0-387-30164-8_576

44. Das A. Logistic Regression. In: Michalos AC, ed. Encyclopedia of Quality of Life and Well-Being Research. Springer Netherlands; 2014:3680–3682. doi:10.1007/978-94-007-0753-5_1689

45. Popescu MC, Balas VE, Perescu-Popescu L, Mastorakis N. Multilayer perceptron and neural networks. WSEAS Trans Cir and Sys. 2009;8(7):579–588.

46. Murtagh F. Multilayer perceptrons for classification and regression. Neurocomputing. 1991;2(5):183–197. doi:10.1016/0925-2312(91)90023-5

47. Guo G, Wang H, Bell D, Bi Y, Greer K. KNN Model-Based Approach in Classification. In: Meersman R, Tari Z, Schmidt DC, eds. On The Move to Meaningful Internet Systems 2003: CoopIS, DOA, and ODBASE. Springer; 2003:986–996. doi:10.1007/978-3-540-39964-3_62

48. Pidsley R, Y Wong CC, Volta M, Lunnon K, Mill J, Schalkwyk LC. A data-driven approach to preprocessing Illumina 450K methylation array data. BMC Genomics. 2013;14(1):293. doi:10.1186/1471-2164-14-293

49. Omer Fadl Elssied N, Ibrahim O, Hamza Osman A. A Novel Feature Selection Based on One-Way ANOVA F-Test for E-Mail Spam Classification. RJASET. 2014;7(3):625–638. doi:10.19026/rjaset.7.299

50. Todorovski L, Džeroski S. Combining Classifiers with Meta Decision Trees. Machine Learning. 2003;50(3):223–249. doi:10.1023/A:1021709817809

51. Wong TT. Performance evaluation of classification algorithms by *k*-fold and leave-one-out cross validation. Pattern Recognition. 2015;48(9):2839–2846. doi:10.1016/j.patcog.2015.03.009

52. Lundberg SM, Nair B, Vavilala MS, et al. Explainable machine-learning predictions for the prevention of hypoxaemia during surgery. Nat Biomed Eng. 2018;2(10):749–760. doi:10.1038/s41551-018-0304-0

53. Ouyang D, Liang Y, Li L, et al. Integration of multi-omics data using adaptive graph learning and attention mechanism for patient classification and biomarker identification. Computers in Biology and Medicine. 2023;164:107303. doi:10.1016/j.compbiomed.2023.107303

54. Mallick H, Porwal A, Saha S, Basak P, Svetnik V, Paul E. An integrated Bayesian framework for multi-omics prediction and classification. Statistics in Medicine. 2024;43(5):983–1002. doi:10.1002/sim.9953

55. Wang KW, Yuan YX, Zhu B, et al. X chromosome-wide association study of quantitative biomarkers from the Alzheimer’s Disease Neuroimaging Initiative study. Front Aging Neurosci. 2023;15:1277731. doi:10.3389/fnagi.2023.1277731

56. Xiong X, James BT, Boix CA, et al. Epigenomic dissection of Alzheimer’s disease pinpoints causal variants and reveals epigenome erosion. Cell. 2023;186(20):4422–4437.e21. doi:10.1016/j.cell.2023.08.040

57. Liu P, Li L, He F, et al. Identification of Candidate Biomarkers of Alzheimer’s Disease via Multiplex Cerebrospinal Fluid and Serum Proteomics. Int J Mol Sci. 2023;24(18):14225. doi:10.3390/ijms241814225

58. Butterfield DA, Boyd-Kimball D, Castegna A. Proteomics in Alzheimer’s disease: insights into potential mechanisms of neurodegeneration. Journal of Neurochemistry. 2003;86(6):1313–1327. doi:10.1046/j.1471-4159.2003.01948.x

