## Supplemental FigureS1-3; Supplemental TableS1-2 for "WIMOAD: Weighted Integration of Multi-Omics Data with Meta Learning for Alzheimer’s Disease Diagnosis": Supplementary_figures.pdf

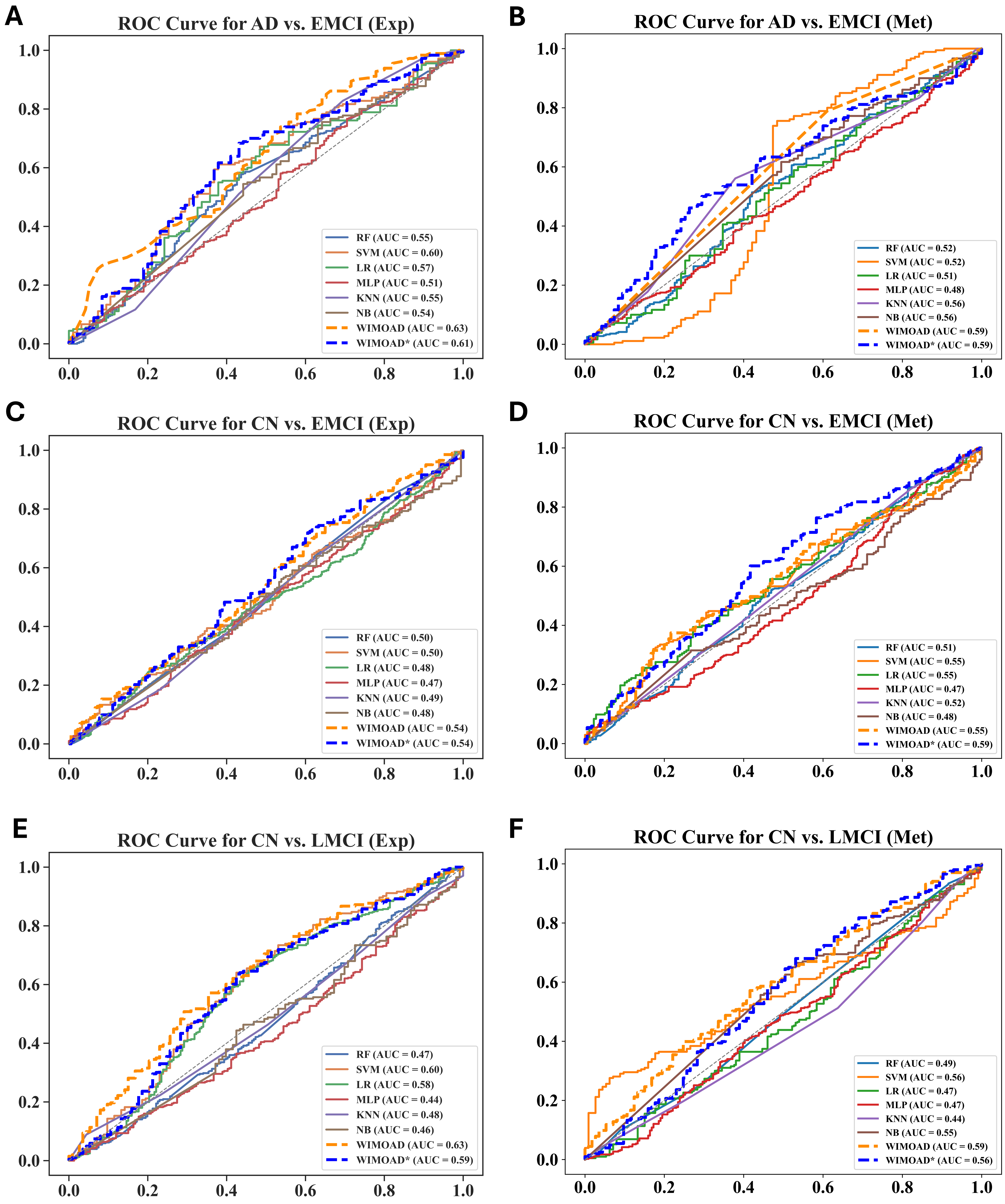


***Supplementary Figure S1. ROC Curve Comparing the Performance of Two Omics using One Classifier and The proposed Model.*** *All classifiers are trained with the same feature dimensions under LOOCV. The performance was measured using the metric introduced previously.* ***(A, C, E)*** *ROC curve on AUC comparison on gene expression data.* ***(B, D, F)*** *ROC curve on AUC comparison on gene methylation data. SVM: Support Vector Machine LR: Logistic Regression; MLP: Multilayer Perceptron; RF: Random Forest; NB: Naïve Bayes; KNN: K-Nearest Neighbor. WIMOAD*: Stacking strategy that adds the selected features for meta model training.*

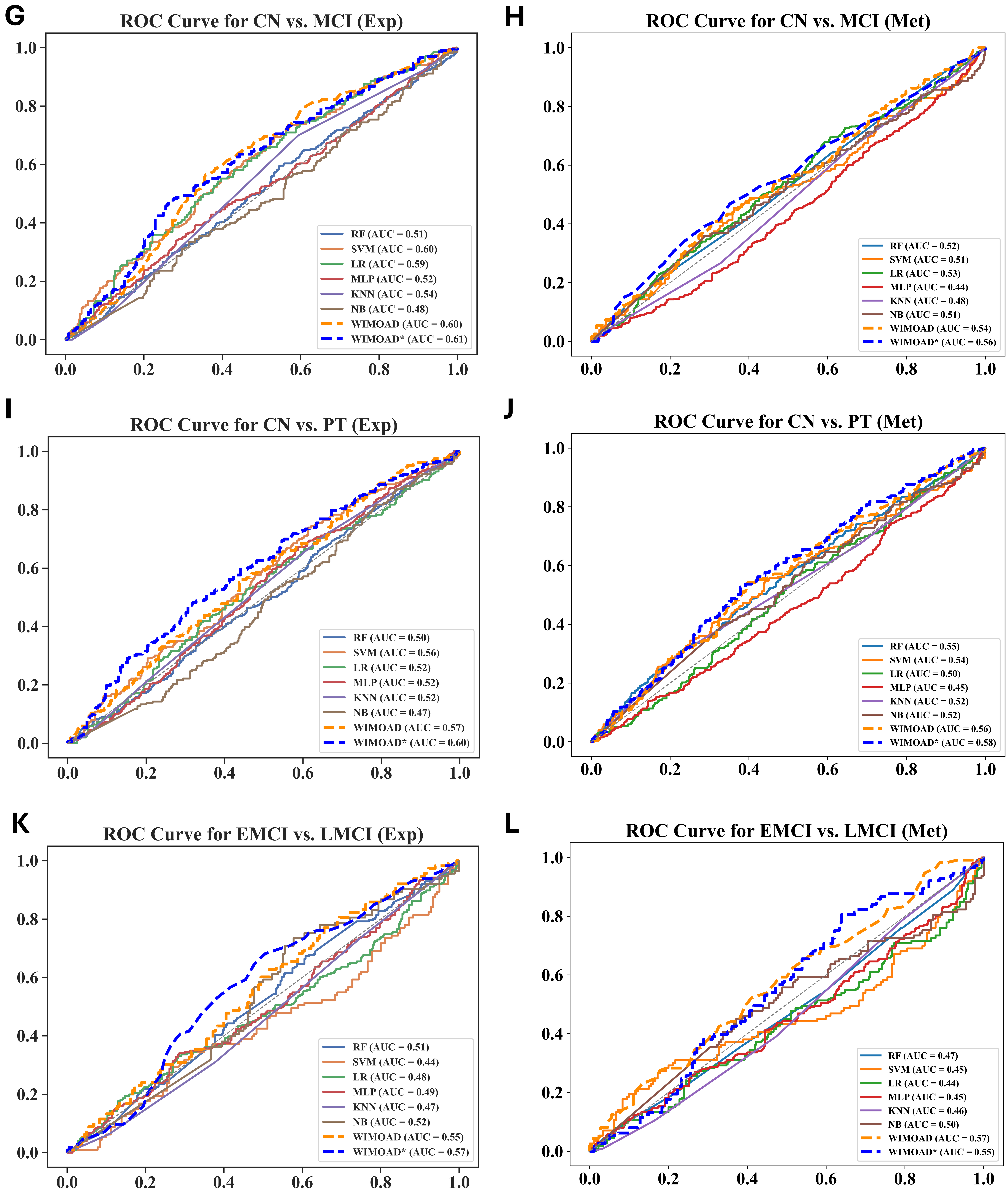


***Supplementary Figure S1 (continued). ROC Curve Comparing the Performance of Two Omics using One Classifier and The proposed Model.*** *All classifiers are trained with the same feature dimensions under LOOCV. The performance was measured using the metric introduced previously.* ***(G, I, K)*** *ROC curve on AUC comparison on gene expression data.* ***(H, J, L)*** *ROC curve on AUC comparison on gene methylation data. SVM: Support Vector Machine LR: Logistic Regression; MLP: Multilayer Perceptron; RF: Random Forest; NB: Naïve Bayes; KNN: K-Nearest Neighbor. WIMOAD*: Stacking strategy that adds the selected features for meta model training.*

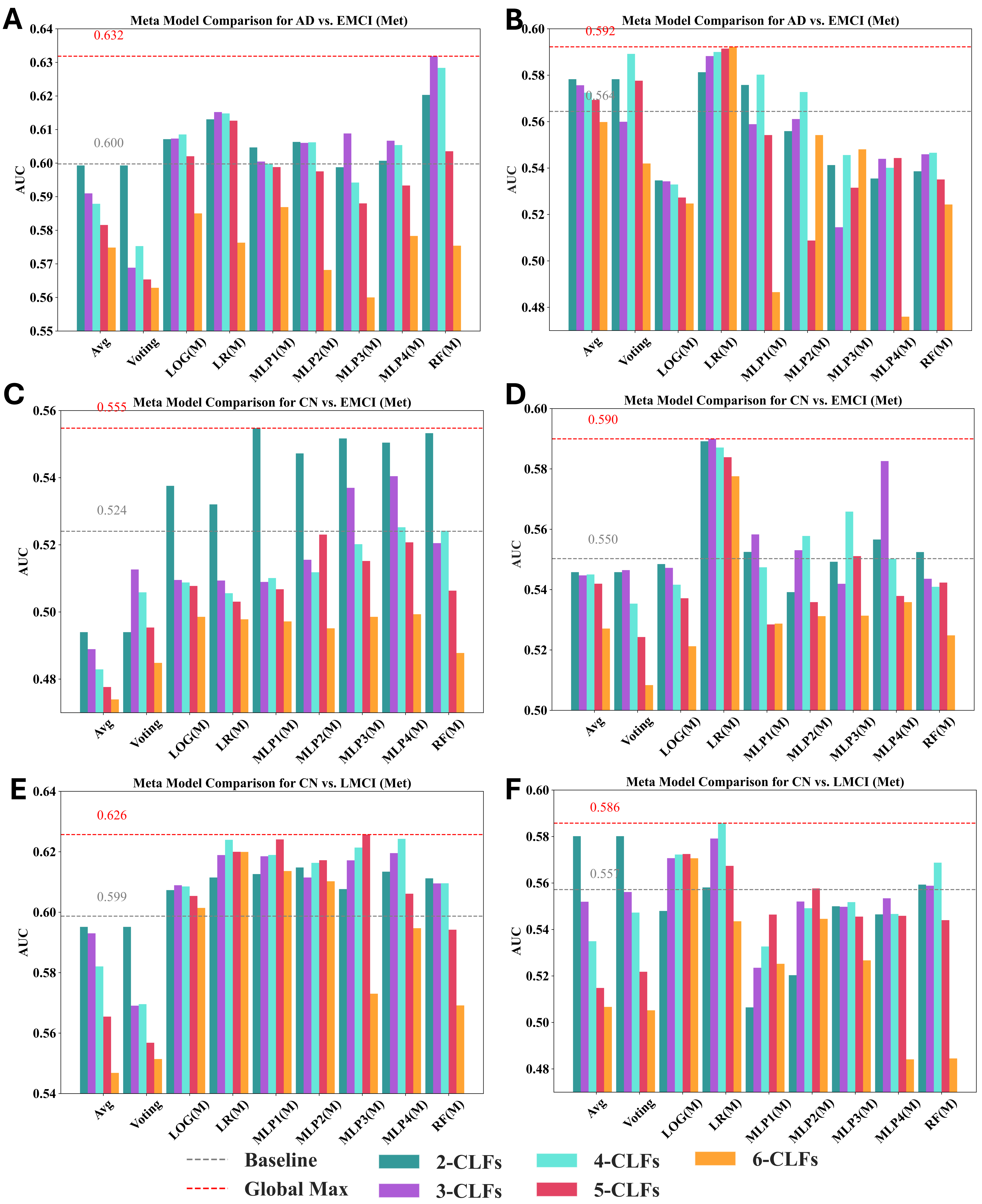


***Supplementary Figure S2. Impact of Meta-Model Selection on AUC Performance for the Same Base Model Combinations. (A, C, E)*** *WIMOAD performance improvement using gene expression data only under different meta models.* ***(B, D, F)*** *WIMOAD performance improvement using gene methylation data only under different meta models. (2 - 6)-CLFs: Number of classifiers used as base models. Avg.: Average the prediction scores of base models for the ensemble; Voting: hard voting strategy for ensemble learning. LOG(M): Logistic Regression as the meta model for stacking ensemble; LR(M): Linear Regression as the meta model for stacking ensemble; MLP(1-4)(M): Multi-Layer Perceptron with different number of hidden layers as the meta model for stacking ensemble; RF(M): Random Forest as the meta model for stacking ensemble. Baseline: The highest AUC score of single classifier. Global Max: the highest AUC score across all combinations of classifiers and meta models in WIMOAD.*

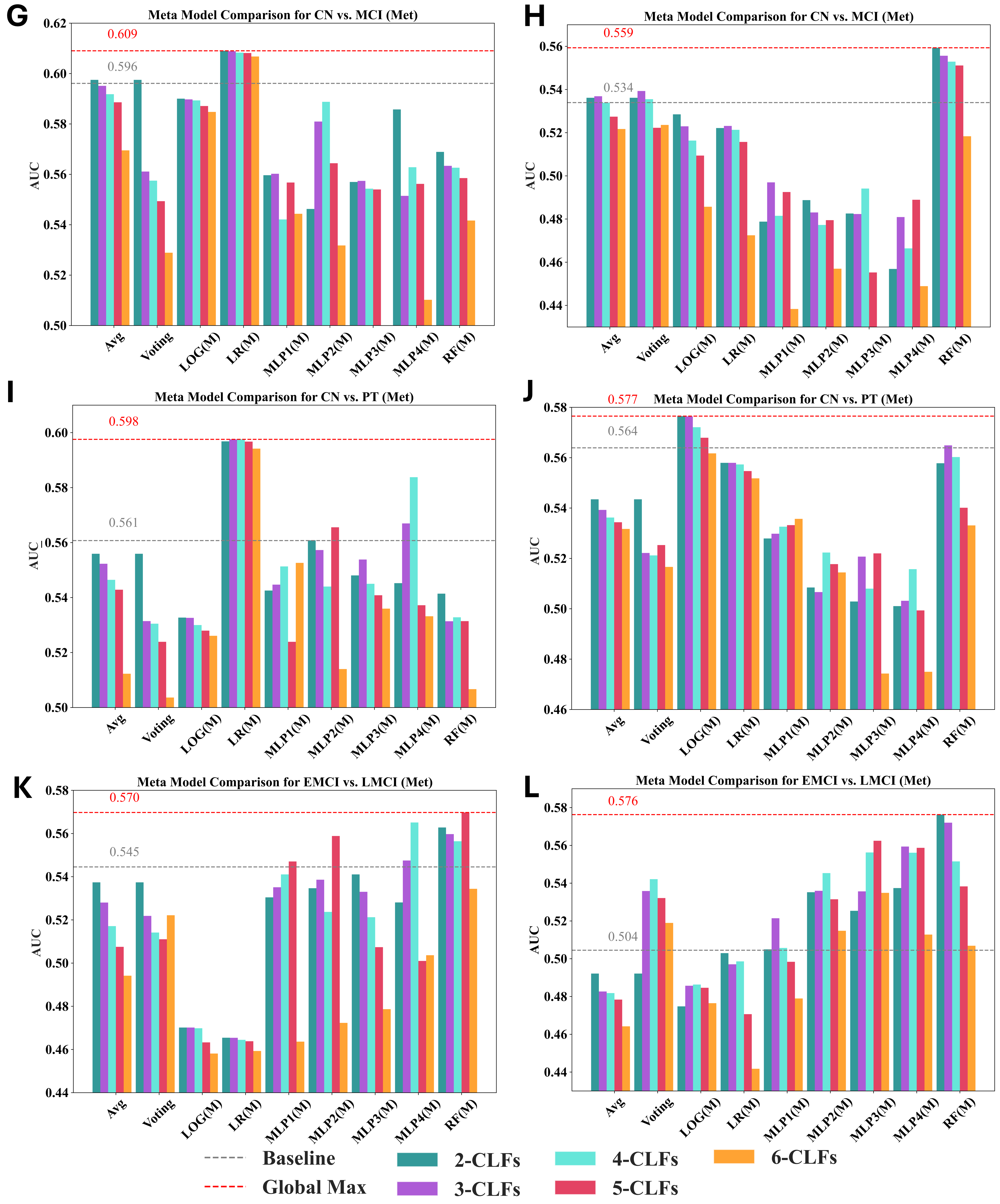


***Supplementary Figure S2 (continued). Impact of Meta-Model Selection on AUC Performance for the Same Base Model Combinations. (G, I, K)*** *WIMOAD performance improvement using gene expression data only under different meta models.* ***(H, J, L)*** *WIMOAD performance improvement using gene methylation data only under different meta models. (2 - 6)-CLFs: Number of classifiers used as base models. Avg.: Average the prediction scores of base models for the ensemble; Voting: hard voting strategy for ensemble learning. LOG(M): Logistic Regression as the meta model for stacking ensemble; LR(M): Linear Regression as the meta model for stacking ensemble; MLP(1-4)(M): Multi-Layer Perceptron with different number of hidden layers as the meta model for stacking ensemble; RF(M): Random Forest as the meta model for stacking ensemble. Baseline: The highest AUC score of single classifier. Global Max: the highest AUC score across all combinations of classifiers and meta models in WIMOAD.*

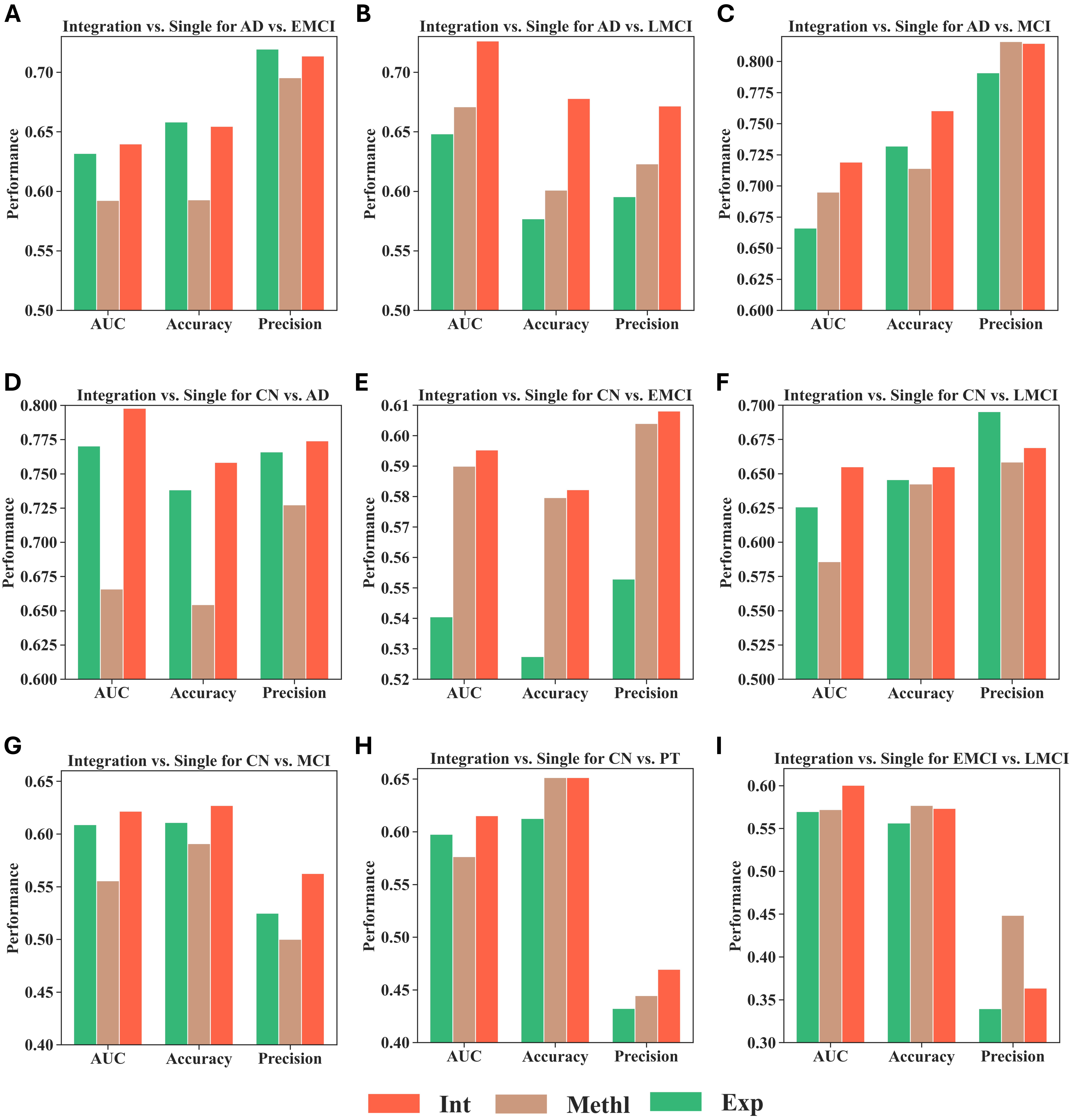


***Supplementary Figure S3 Integration Performances of WIMOAD.*** *The x-axis represents the evaluation matrics, and the y-axis represents performance of the models. The results were generated under the best coefficient selected. The Integration model improvement on the prediction is statistically significant according to the McNemar's Test.*
